# Detection of Potential Phytochemicals against ctxAB Toxin to Combat Cholera

**DOI:** 10.1101/2024.07.29.605560

**Authors:** Yasmin Akter, Khalilur Rahman, Md Selim Reza, Abdullah Al-Mamun, Munirul Alam, Md Nurul Haque Mollah, Md. Mamun Monir

## Abstract

*Vibrio cholerae* is a gram-negative curved rod-shaped bacterium responsible for cholera, an intestinal infection characterized by severe acute watery diarrhea that can be fatal if untreated. The major virulence factor for *Vibrio cholerae* is cholera toxin (*ctx*), a potent toxin which has two subunits, A and B (ctxAB), that are crucial for the progression of the deadly disease. The B subunit is a pentavalent protein that binds to the intestinal mucosa to allow internalization of the A subunit to initiate rice-watery diarrhea. This study explored potential phytochemicals to inhibit the B subunit from binding to the host mucosa by examining their interaction with *ctxB*. Proteins encoded by the three genotypes of the *ctxB* gene (*ctxB1, ctxB3*, and *ctxB7*) present in the El Tor biotype strains of *V. cholerae* O1 responsible for the currently ongoing seventh cholera pandemic were targeted. Analysis of 52 phytochemicals obtained from PubChem identified a group of phytochemicals based on their binding affinity scores with the proteins of *ctxB* genotypes. Assessment of drug-likeliness and toxicity risk highlighted two phytochemicals, Limonin and Emodin, as the best potential candidates for inhibiting the mode of action of cholera toxin. Molecular dynamic (MD) simulation-based MM-PBSA analysis suggested these compounds stably interact with the receptors. Molecular docking and molecular dynamic simulations revealed that these phytochemicals might have potential to inhibit the cholera toxin function and reduce the severity of cholera.

**Author Summary:** Cholera toxin (ctx) subunits A and B (ctxAB) play a significant role in the disease, with the B subunit facilitating the binding of the toxin to the intestinal cells, and the A subunit entering the cells, resulting in the cellular changes that lead to watery diarrhea. To inhibit the B subunit of cholera toxin from binding host mucosa, we explored candidate phytochemicals instead of synthetic molecules as drug agents by examining their binding behavior with ctxB using a molecular docking approach. The molecular docking, molecular dynamic simulations, and toxicity analyses identified two phytochemical compounds that have the potential to inhibit the mechanism of cholera toxin.

## Introduction

Cholera is an acute form of secretory diarrhea, caused by the gram-negative proteobacterium, *Vibrio cholerae*. The clinical manifestations of cholera involve the continual discharge of abundant fluid exhibiting a rice-water consistency, often leading to severe dehydration [1]. As per the reports of WHO, 2.9 million cases of cholera emerge every year in the approximately 69 cholera-endemic countries and the estimated number of deaths is around 21,000 to 1,43,000 per year [2]. Seven cholera pandemics have been recorded in history since 1817, and the current seventh pandemic started in the early 1960s due to the emergence of *Vibrio cholerae* O1 biotype El Tor; while the previous pandemics were linked to the classical biotype of the bacterium [3]. El Tor strains are superior to classical strains in terms of host-to-host transmission efficiency and environmental and human host viability [4]. The El Tor strain exhibiting increased gene expression associated with biofilm formation, chemotaxis, and the transfer of iron, peptides, and amino acids, might be a potential reason for the recent upsurge of El Tor cholera worldwide due to the elevated ability of the bacterium to thriving in the aquatic environmental reservoirs [5].

The first-line treatment regimen for cholera is the oral rehydration saline (ORS) [6] fluids. Although ORS is considered the best therapeutic measure, its efficacy is limited. This is because the administration of ORS alone does not alleviate symptoms, eradicate *V. cholerae* infection, or disrupt the mechanism of action of cholera toxin (CT) [7]. Besides, two doses of effective antibiotics are a convention to shorten the duration of hospitalization and reduce the burden of the pathogen to prevent further disease transmission. Antibiotics like tetracycline, azithromycin, erythromycin, and fluoroquinolone have been employed for cholera disease treatment over the years [8-11]. Unfortunately, extensive use of these antibiotics has demonstrated a concerning trend of developing resistance or multidrug resistance, which is attributed to the acquisition of mobile genetic elements or mutations in one or more genes [12,13]. Bacteriophage therapy is also a potential tool to use against *V. cholerae* to combat cholera diseases but it has certain drawbacks because bacteria can make themselves resistant to phage infections by inhibiting phage genome injection through the super injection exclusion (Sie) mechanism, a protein-mediated process expressed by prophage [14].

In the course of infection, the cholera toxin (CT) takes part in binding with the specific receptor binding site, the enterocyte monosialoganglioside GM1, on host enterocytes which results in increased levels of cAMP levels causing major secretion of profuse diarrhea filled with water and electrolytes [15]. When the cholera toxin attaches itself with high precision and affinity to the receptor residing on the GM1, there occurs an increase in intestinal secretion, initiating endocytosis and instigating a sustained impact on the enterocyte cytosol [16]. Thus, we designed a therapeutic strategy in which the binding of the cholera toxin with monosialoganglioside GM1 can be prevented with phytochemicals without any adverse effects, which is very crucial. In this study, we conducted molecular docking for identifying potential phytochemicals instead of synthetic molecules as drug molecules that can inhibit cholera toxin. The phytochemicals are preferred over synthetic drugs because the latter might have some adverse effects on the human body [17,18]. For this, we tested 52 phytochemical compounds from PubChem for binding affinity and also examined the drug ability and toxigenicity of the molecules. Although numerous phytochemicals are accessible from PubChem, 52 phytochemicals used in the current study have been selected based on their high antimicrobial, antiviral, and antidiarrheal properties [19-21]. The selected phytochemicals generally display their antimicrobial properties by interfering with the intermediary mechanism, starting the process of cytoplasmic components, altering DNA/RNA synthesis and function, and interfering with regular cell communication [22].

## Materials and Methods

### Target proteins encoded by the genotypes of the *ctxB* gene

Initially, the El Tor strains were characterized by the *ctxB3* genotype. Since then, they have seen evolutionary modifications and have acquired the classical cholera toxin genotype 1 (*ctxB1*) [23]. The Haitian cholera toxin (Haitian CT) is a novel variation of the *V. cholerae* O1 El Tor biotype that appeared after the devastating cyclone in Odisha in October 1999 [2], and in November 1999, the Erasama block in the Jagatsinghpur district reported the very first case of cholera with the *ctxB7* genotype [23]. In this study, we considered proteins encoded by the three genotypes *ctxB1, ctxB3*, and *ctxB7* of the El Tor biotype of *V. cholerae* O1 for molecular docking and dynamic simulation.

### Molecular docking and pharmacokinetics analyses

The amino acid sequence of *ctxB* genotypes collected from NCBI was used for constructing 3D structures using SWISS-MODEL [24]. The 3D structures of the target proteins were visualized using Discovery Studio Visualizer 2019 [25]. PDB2PQR and H++ were used to determine the protonation status of the proteins [26,27]. After that, water molecules, attaching ligands, and heteroatoms were removed using AutoDock tools [28,29]. In addition, the software was used for setting the torsion tree, rotatable and nonrotatable bonds in the phytochemicals, and adding important hydrogen atoms to the target proteins, particularly in the site of binding, though these proteins can be associated with any kind of interaction with the ligands. Then, the entire protein surface was covered by a grid box.

The 3D structures of the phytochemical compounds were downloaded from the PubChem database, and Avogadro was utilized to minimize the energy of the phytochemicals [30,31]. Then, the binding affinities for each of the target proteins and phytochemical compounds were calculated using AutoDock Vina [32]. PyMol [33] and Discovery Studio Visualizer 2019 [25] were used for visualizing the surfaces of complexes, non-covalent interaction types, and distances.

In order to select top-ranked targets and agents, let *X*_*pq*_ represents the binding affinity score corresponding to the *p*^th^ target protein (*p* = 1, 2, … *r*) and *q*^th^ phytochemicals (*q* = 1, 2, … *s*). Then, descending row sums, 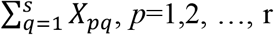, sorted target proteins as the receptors and column sums, 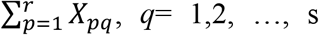, sorted lead phytochemicals as the candidate drug agent. Additionally, the canonical SMILES of the top-ranked phytochemicals were used as input for the pharmacokinetic assessments in two integrated online tools-SwissADME [34], and pkCSM [35], that precisely predict the phytochemical pharmacokinetics, drug-like characteristics, and pharmaceutical chemistry-friendly properties.

### Molecular dynamic (MD) simulations

The molecular dynamics simulations of the chosen compounds with an AMBER14 force field [36,37] were performed using the YASARA software [38] to ascertain how the top-ranked protein-phytochemical complexes behave dynamically. The MD simulation was performed using nine distinct complexes, comprising three variants of *ctxB* (*ctxB1, ctxB3*, and *ctxB7*), each of which interacted with dehydropipernonaline, emodin, and limonin. The hydrogen bonding structure of the protein-phytochemical complexes was modified and solved with a TIP3P water model. The hydrogen bonding network of the protein-phytochemical complexes was altered and solved using a TIP3P water model before the simulation [39], and 0.990 gL^-1^ solvent density was used to sustain periodic boundary conditions. During solvation, pKa calculations were performed on the complexes containing titratable amino acids, and at 298 K, pH 7.4, and 0.9 percent NaCl, the simulated systems reached neutralization [40]. The initial energy minimization procedure for all simulation systems, consisting of 23,358, 23,963, and 23,663 atoms for the *ctxB1*-Dehydropipernonaline, *ctxB1*-Emodin, and *ctxB1*-Limonin complexes, respectively, and 23,263, 24,157, and 24,062 atoms for the *ctxB3*-Dehydropipernonaline, *ctxB3*-Emodin, and *ctxB3*-Limonin complexes, and 23,379, 23,707, and 25,038 atoms for the *ctxB7*-Dehydropipernonaline, *ctxB7*-Emodin, and *ctxB7*-Limonin complexes, respectively. The steepest gradient method was used for this optimization process, which involved 5000 cycles for each complex. Every 250 ps, the trajectories were recorded to carry out the subsequent analysis using macro and SciDAVis software (available at http://scidavis.sourceforge.net/). Then, for each snapshot, the MM-Poisson-Boltzmann surface area (MM-PBSA) binding free energy calculation was performed using the following formula in the YASARA script [41,42].

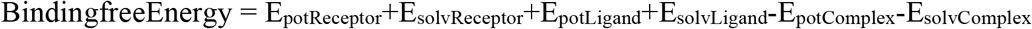

Here, AMBER-14 and the built-in YASARA macros were used as the force field to calculate the MM-PBSA binding energy; higher positive energies suggest improved binding [43].

## Results

### 3D structure of the mutants of *ctxB*

*Vibrio cholerae* carries the CT prophage gene, responsible for encoding the cholera toxin which leads to clinical symptoms of the deadly disease cholera [44]. 12 genotypes of *ctxB* toxin have been identified so far, of which three (*ctxB3, ctxB1, and ctxB7*) can be identified from the seventh pandemic El Tor strains. Recent genomic analyses showed *V. cholerae* O1 strains circulating at the Asian (Asian lineage) region contain *ctxB1*, the strains circulating globally (global lineage) contain *ctxB7*, and *ctxB3* genotype strains seem to exist no longer [13]. Three-point mutations differentiated the three genotypes of *ctxB*, where both *ctxB1* and *ctxB7* commonly have 2 mutations and the other one has an additional point mutation (**Fig 1a**). 3D structures of the proteins encoded by three different genotypes of *ctxB* showed that *ctxB1* and *ctxB7* encoded proteins had different configuration than the *ctxB3* encoded protein, which might have influence on binding affinities with the ligands (**Fig 1b-d**).

**Fig 1.**
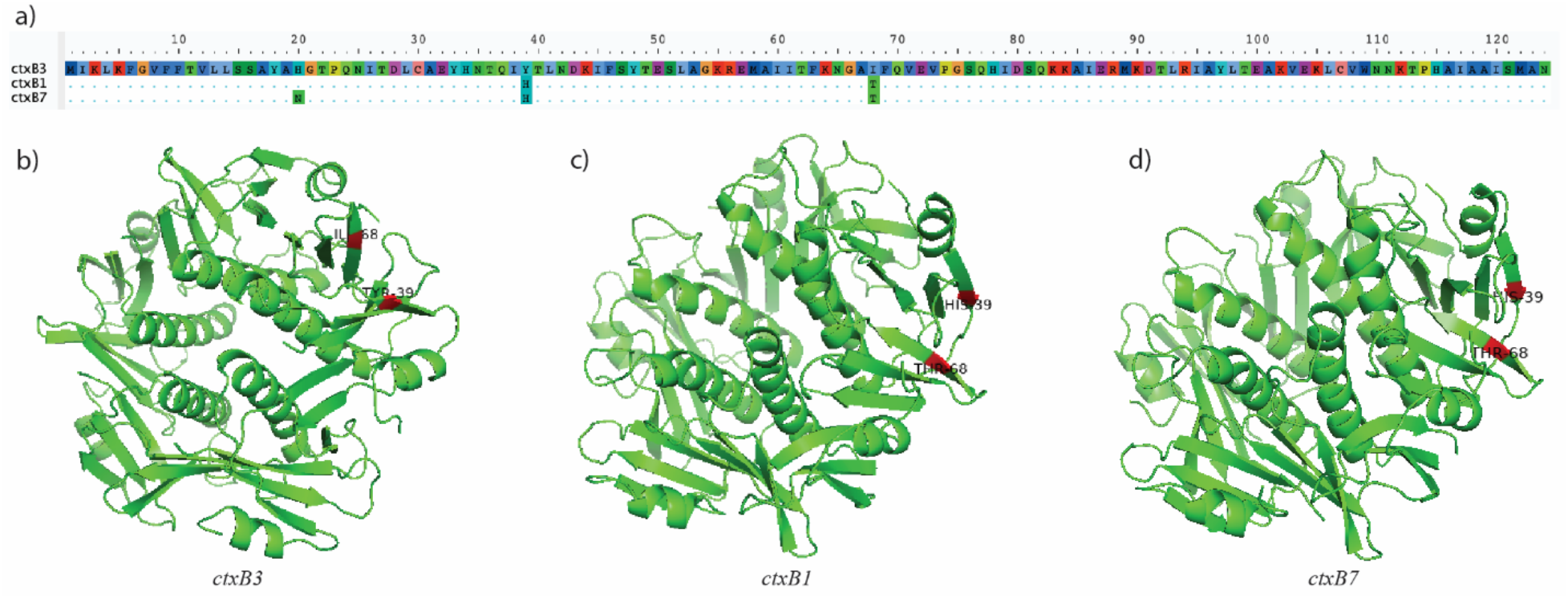
*ctxB* genotypes, binding sites, and their 3D structures. (a) shows the amino acid sequences of the three mutated genotypes of cholera B toxin. *ctxB1* shows amino acid substitution at positions 39, and 68, while *ctxB7* undergoes additional mutation at the 20^th^ position [45]. (b-d) displays the 3D structures modeled from the amino acid sequences of the three genotypic variants of the *ctxB* toxin. The 3D models were visualized using the Pymol [46] software and the area of mutations have been highlighted in red.

### Drug Repurposing by Molecular Docking

Molecular docking was carried out for the 52 phytochemicals against encoded proteins of *ctxB1, ctxB3*, and *ctxB7*. A total of r = 3 receptors and s = 52 phytochemical compounds were used in the molecular docking process to determine the binding affinity scores (kcal/mol) for each pair of phytochemical compounds and receptors. Afterward, using the row sums of the binding affinity matrix **X** = (X_pq_) to sort the target receptors and the column sums of the phytochemical compounds, we were able to identify just a small number of these compounds as potential candidates (**Fig 2**). The top 10 phytochemical agents were Limonin, Corilagin, Taraxesterol, Procyanidin, Beta-carotene, Emodin, Dehydropipernonaline, Apigenin, Guineesine, and Benzethonium based on the analysis results with binding affinity scores against the three receptor proteins encoded by *ctxB1, ctxB3*, and *ctxB7* **(Table S1 in supplementary information**).

**Fig 2.**
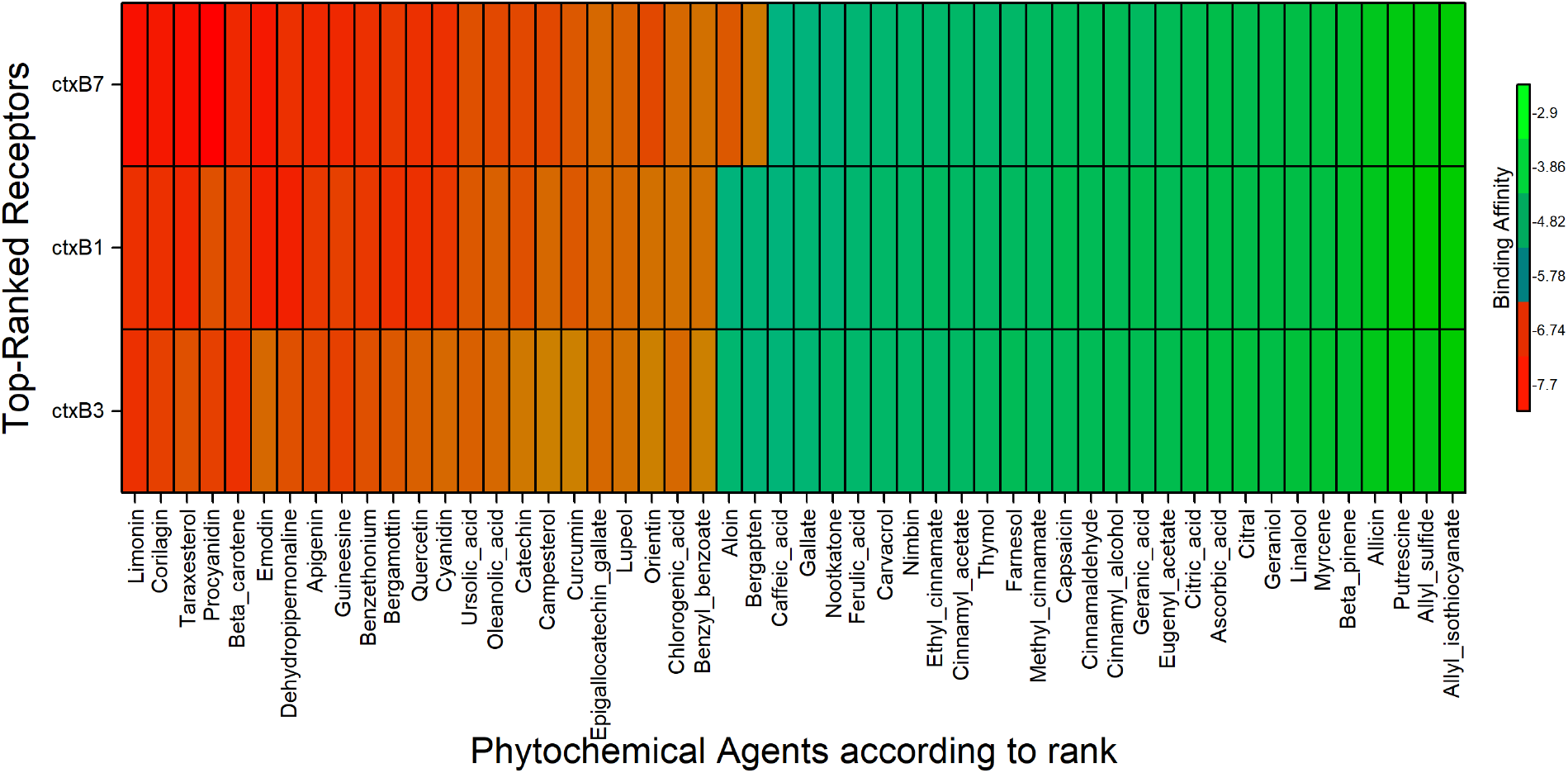
Binding affinities of the top-ordered phytochemical agents against the 3 receptors. Here, orange to red colors indicate the higher binding affinities of the phytochemicals. A total of 23 phytochemicals were identified with moderate to high binding affinities (<= -6.3) for *ctxB1* and *ctxB3*. Two additional phytochemicals had moderate binding affinities for *ctxB7*.

### Pharmacokinetics of the best phytochemicals

The ADME (Absorption, Distribution, Metabolism, and Excretion) properties of phytochemicals were determined using the ADME predictor [34,35]. The toxicity risk, drug store viability, and drug-likeness of each screened phytochemical were examined. The degree of indemnity and efficacy of the top three lead medications were established by pharmacokinetic characteristic analysis (**Table 1**). The significant factors that may influence the top-ranked phytochemicals were taken into account, such as molar refractivity, lipophilicity, water solubility, GI absorption, CNS permeability, P-gp substrate, bioavailability score, skin permeation, and Lipinski condition. The analyses revealed three phytochemicals, dehydropipernonaline, emodin, and limonin, as the best candidates for additional screening. They all meet the requirements to be considered as drugs, including lipophilicity (logP <5), molar refractivity of 40-130, high GI absorption, and water solubility (logS >-5). Our chosen phytochemicals comply with Lipinski’s rule, which states that orally active medications generally show no more than one violation of particular physicochemical characteristics, suggesting a good chance of oral absorption and efficacy. Finally, we verified whether our suggested phytochemicals were toxic or not for humans or rats. We find dehydropipernonaline with hepatotoxicity (**Table S2 in supplementary information**).

**Table 1.**
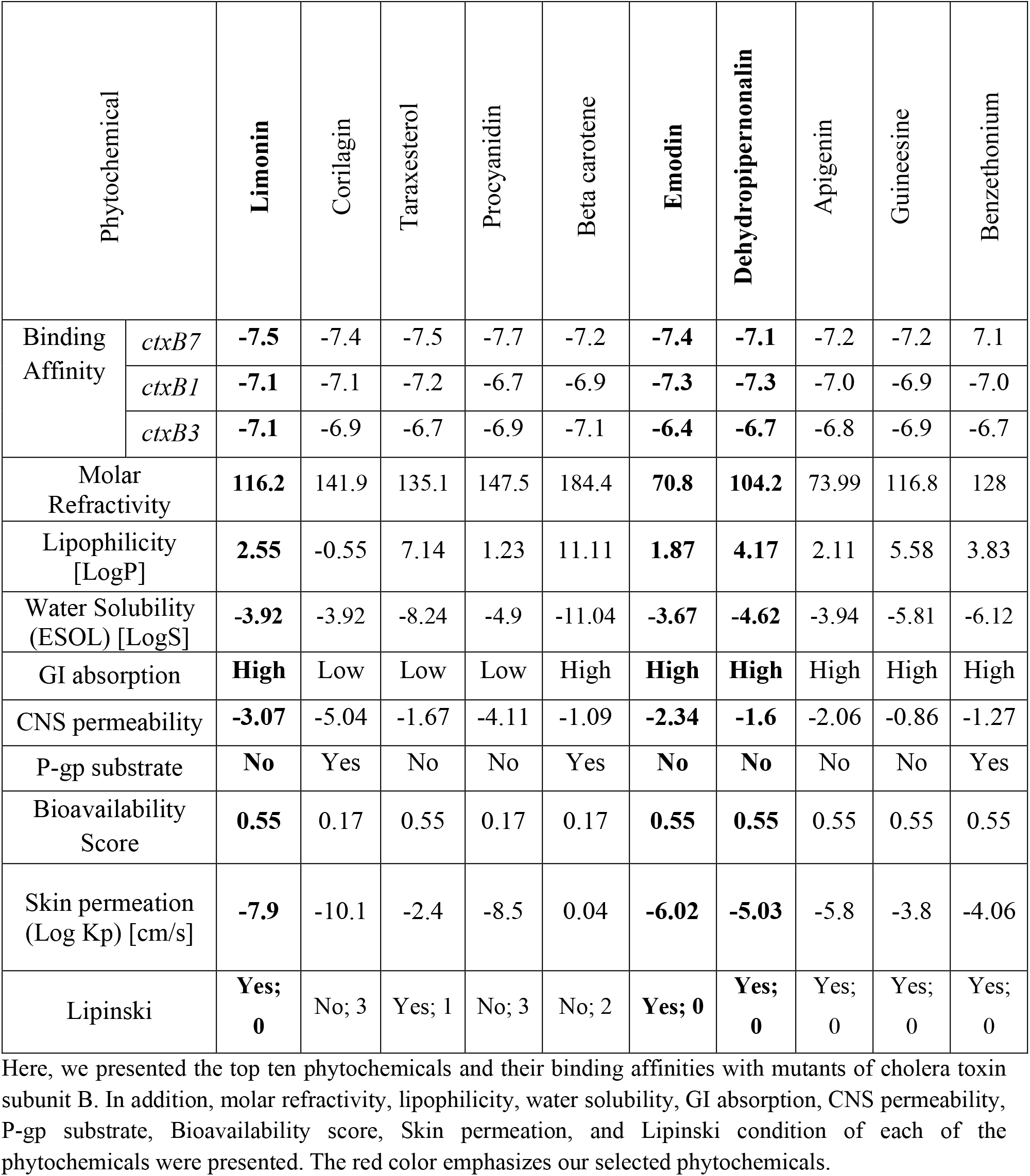
Pharmacokinetics of the best phytochemicals.

### Binding sites of *ctxB* and phytochemicals

The GM1 pentasaccharide receptor binds with the protein encoded by the wild-type *ctxB* (classical, similar to *ctxB1*) at residues Asn111, Asn35, Lys112, Glu72, Gln82, His78, Ile79, Glu32, Trp109, Tyr33, and His34 [47]. It is therefore interesting to examine the binding sites of phytochemicals with proteins of *ctxB* genotypes to understand their potential inhibitory properties. To explore this, we analyzed the docked ligand-protein complexes and visualized their 2D and 3D structures (**Fig 3**). The analysis revealed that the *ctxB1*-Dehydropipernonaline complex forms two hydrogen bonds and six other hydrophobic bonds. The hydrogen bonds occur at residues Lys112 (conventional) and Asn111 (carbon), while the hydrophobic bonds are at Trp109 (pi-pi stacked and pi-alkyl), Ala85 (alkyl), Ile86 (alkyl), Met89 (alkyl), and Tyr33 (pi-alkyl) (**Table 2**). Moreover, dehydropipernonaline and GM1 share four common binding amino acid positions to bind with the *ctxB1* protein. The *ctxB1*-Emodin complex forms three conventional hydrogen bonds at residues Gln82, Lys112, and Asn111, along with three hydrophobic bonds at Trp109 (pi-stacked) and Tyr33 (pi-pi T-shaped and pi-alkyl). Consequently, emodin and GM1 share five common amino acid binding positions to bind with *ctxB1*. The *ctxB1*-Limonin complex forms two hydrogen bonds at residues Gln82 (conventional) and Ala118 (carbon), and one hydrophobic bond at Ile86 (pi-alkyl). Thus, limonin and GM1 share only one common amino acid position to bind with *ctxB1*. Structure of the protein encoded by *ctxB7* is pretty similar to *ctxB1* (**Fig 1c-d**). Hence, the binding positions of *ctxB7*-Dehydropipernonaline complex are similar to the *ctxB1*-Dehydropipernonaline except for an additional hydrogen bond at the Ala85 (carbon) residue. The amino acid residue Lys112 of *ctxB1*-Emodin complex doesn’t exist to forming the *ctxB7-*Emodin complex. Lastly, the formation of the *ctxB7-*Limonin complex is pretty different from *ctxB1-*Limonin. This complex (*ctxB7-*Limonin) is formed by two hydrogen bonds at residues Lys112 (conventional) and Glu72 (carbon), and one hydrophobic bond at Trp109 (pi-pi stacked), and all three positions are common for Limonin and GM1 to bind with protein encoded by *ctxB7*.

**Table 2.** Non-bond interactions between three top-ordered receptors and phytochemicals.

| Phytochemical/<br>ligand | Receptor/<br>Target | Interacting<br>Amino Acid | Interaction<br>Category | Interaction<br>Type |
| --- | --- | --- | --- | --- |
| Dehydropipernonaline | *ctxB1* | Lys112 | Hydrogen | Conventional |
|  |  | Asn111 | Hydrogen | Carbon |
|  |  | Trp109 | Hydrophobic | Pi-Pi Stacked |
|  |  | Ala85 | Hydrophobic | Alkyl |
|  |  | Ile86 | Hydrophobic | Alkyl |
|  |  | Met89 | Hydrophobic | Alkyl |
|  |  | Tyr33 | Hydrophobic | Pi-Alkyl |
|  |  | Trp109 | Hydrophobic | Pi-Alkyl |
| Emodin | *ctxB1* | Gln82 | Hydrogen | Conventional |
|  |  | Lys112 | Hydrogen | Conventional |
|  |  | Asn111 | Hydrogen | Conventional |
|  |  | Trp109 | Hydrophobic | Pi-Pi Stacked |
|  |  | Tyr33 | Hydrophobic | Pi-Pi T-shaped |
|  |  | Tyr33 | Hydrophobic | Pi-Alkyl |
| Limonin | *ctxB1* | Gln82 | Hydrogen | Conventional |
|  |  | Ala118 | Hydrogen | Carbon |
|  |  | Ile86 | Hydrophobic | Pi-Alkyl |
| Dehydropipernonaline | *ctxB3* | His78 | Hydrogen | Conventional |
|  |  | Leu29 | Hydrophobic | Alkyl |
|  |  | Arg88 | Hydrophobic | Alkyl |
|  |  | Met89 | Hydrophobic | Alkyl |
|  |  | Ala119 | Hydrophobic | Alkyl |
|  |  | Ala85 | Hydrophobic | Pi-Alkyl |
|  |  | Ile86 | Hydrophobic | Pi-Alkyl |
| Emodin | *ctxB3* | Gln82 | Hydrogen | Conventional |
|  |  | His78 | Hydrogen | Carbon |
|  |  | Trp109 | Hydrophobic | Pi-Pi Stacked |
|  |  | Tyr33 | Hydrophobic | Pi-Pi T-shaped |
|  |  | Ile79 | Hydrophobic | Alkyl |
| Limonin | *ctxB3* | Gln82 | Hydrogen | Conventional |
|  |  | Ala118 | Hydrogen | Carbon |
|  |  | Ile86 | Hydrophobic | Pi-Alkyl |
| Dehydropipernonaline | *ctxB7* | Lys112 | Hydrogen | Conventional |
|  |  | Ala85 | Hydrogen | Carbon |
|  |  | Asn111 | Hydrogen | Carbon |
|  |  | Trp109 | Hydrophobic | Pi-Pi Stacked |
|  |  | Ala85 | Hydrophobic | Alkyl |
|  |  | Ile86 | Hydrophobic | Alkyl |
|  |  | Met89 | Hydrophobic | Alkyl |
|  |  | Tyr33 | Hydrophobic | Pi- Alkyl |
|  |  | Trp109 | Hydrophobic | Pi- Alkyl |
| Emodin | *ctxB7* | Gln82 | Hydrogen | Conventional |
|  |  | Asn111 | Hydrogen | Conventional |
|  |  | Trp109 | Hydrophobic | Pi-Pi Stacked |
|  |  | Tyr33 | Hydrophobic | Pi-Pi T shaped |
|  |  | Tyr33 | Hydrophobic | Pi-Alkyl |
| Limonin | *ctxB7* | Lys112 | Hydrogen | Conventional |
|  |  | Glu72 | Hydrogen | Carbon |
|  |  | Trp109 | Hydrophobic | Pi- Pi Stacked |

Here, the term "phytochemical/ligand" refers to the top-ranked phytochemicals that have been determined by their pharmacokinetic and binding affinities; Receptor/target refers to three mutants of cholera toxin subunit B; Interacting amino acid refers to the binding sites of the ligands; Interaction Category refers to the type of chemical bonds between cholera toxin subunit B with phytochemicals; and Interaction Type refers to the specific types of the bonds.

**Fig 3.**
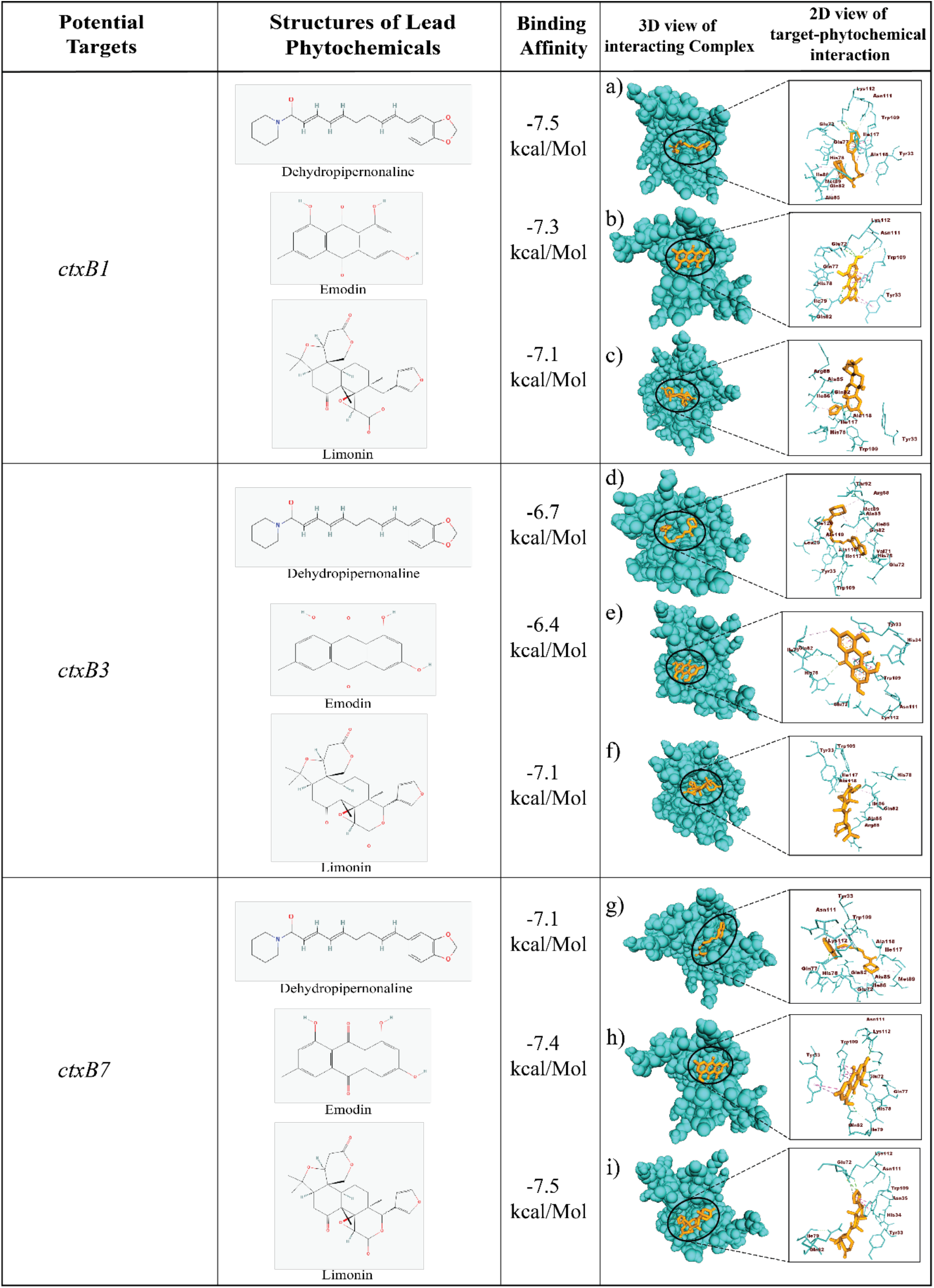
Nine complexes, and their 2D, and 3D chemical interactions. Using Discovery Studio visualizers, the three-dimensional view of interacting complexes was displayed [21]. The complexes are indicated: (a) *ctxB1*-Dehydropipernonaline; (b) *ctxB1*-Emodin; (c) *ctxB1*-Limonin; (d) *ctxB3*-Dehydropipernonaline; (e) *ctxB3*-Emodin; (f) *ctxB3*-Limonin; (g) *ctxB7*-Dehydropipernonaline; (h) *ctxB7*-Emodin; and (i) *ctxB7*-Limonin.

In addition, the complex of *ctxB3* and Dehydropipernonaline forms one conventional hydrogen bond at His68, and six hydrophobic bonds at Leu29, Arg88, Met89, Ala119, Ala85 (pi-alkyl), and Ile86 (pi-alkyl) residues. *ctxB3*-emodin complex forms two hydrogen bonds at Gln82 (conventional), and His78 (carbon) and three hydrophobic bonds at Trp109 (pi-pi stacked), Tyr33 (pi-pi T-shaped), Ile79 (Alkyl) residues. And, *ctxB3*-Limonin complex forms two hydrogen bonds at Gln82 and Ala118, and one hydrophobic bond at Ile86.

### Molecular Dynamic Simulations

In our analysis, limonin, emodin, and dehydropipernonaline were the top three candidate phytochemicals among those that were revealed as the best candidates. Thereby, the top three phytochemical compounds were selected for 100 ns MD-based MM-PBSA simulations to be used for stability study. Every complex, with the exception of the *ctxB3* and emodin complex, was remarkably stable between the initial and moving phytochemical-target complex changes (**Fig 4**). For each of the three selected phytochemicals, we determined the root mean square deviation (RMSD), and MM-Poisson-Boltzmann surface area (MM-PBSA) corresponding to the receptors.

**Fig 4.**
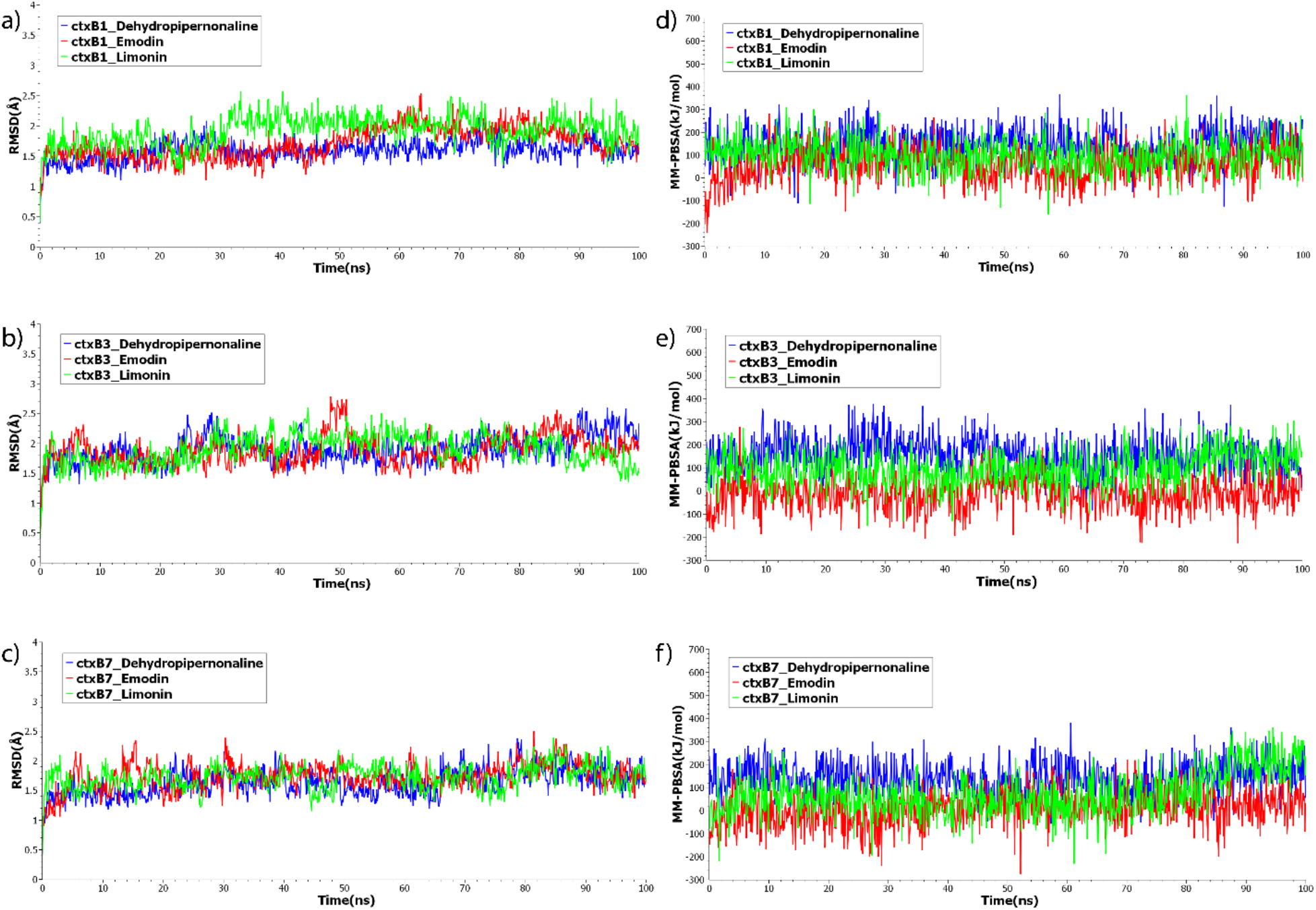
Molecular Dynamic simulation results. The root means square deviations (RMSDs) of the three protein backbone atoms (C, Cα, and N) are displayed in (**Fig 4a-c)** over time for every docked complex. The binding free energy (in kJ mol^−1^) of each snapshot, was calculated and is displayed in (**Fig 4d-f)**. Positive values signify stronger binding. Phytochemicals: Green indicates limonin, red indicates emodin, and blue indicates dehydropipernonaline.

An average RMSD less than 2.5Å is considered binding stability for the complexes [48]. Analyses showed the average RMSD was between 1.59-1.91 Å for all of the nine complexes, which indicates their binding stability. As seen in (**Fig 4a)**, every complex displayed almost stable RMSD and MM-PBSA over the times, except the complexes between *ctxB1* and limonin and *ctxB3* and emodin, which fluctuate frequently during the first few nanoseconds. Eventually, all complexes reached a stable state after 60 ns of the experiment. Here, we determined the MM-PBSA binding free energy for each of the nine complexes that resulted from the pairing of three chosen phytochemicals with three previously identified receptors. The average binding energies were between 58.227 and 142.734 kJ/mol for *ctxB1-*phytochemical complexes, between -15.038 and 157.331 for *ctxB3-*phytochemical complexes, and between 5.837 and 140.373 kJ/mol for *ctxB3-*phytochemical complexes. Remarkably, we observed that *ctxB3* and emodin had average negative binding energies in dynamic simulation, supporting the molecular docking analysis result for this complex **(Fig 3**).

## Discussion and Conclusion

Emergence of multiple antimicrobial resistance against the available antibiotics for treating infectious diseases is a grand challenge which demands alternative therapeutic agents to save lives. In this study, we identified alternative medications for cholera disease, which act by inhibiting the mode of action of cholera toxin. Cholera toxin is a traditional A-B type toxin, which functions constitutively activating AC and causing G protein to be ADP-ribosylated, which raises cyclic AMP levels in the host cell, the rapid discharge of chloride ions by the cystic fibrosis transmembrane conductance regulator (CFTR), the decreased influx of sodium ions, and massive water release via intestinal cells causes acute diarrhea, and vomiting, indicating the initial manifestations of cholera [1]. The cholera toxin initiates its pathogenic effect through the high-affinity attachment of its B subunits to the monosialoganglioside GM1 receptors of the host [49]. Since the key toxin responsible for the pathogenicity of *V. cholerae* is the *ctxB*, inhibiting this gene function may prevent the cholera bacterium from acting in its intended manner. We employed a molecular docking technique to find potential phytochemicals as drug agents with antibacterial properties since phytochemicals have lower adverse effects compared to the synthetic drugs [17,18]. We investigated potential phytochemicals by assessing their binding behavior against the CTX subunit B using a molecular docking technique to prevent the B subunit from binding with host mucosa. Molecular docking was used to target proteins encoded by three genotypes of the *ctxB* gene (*ctxB1, ctxB3*, and *ctxB7*) of the El Tor biotype of *Vibrio cholerae* O1, which is responsible for the currently ongoing 7th cholera pandemic. We examined a pool of 52 phytochemicals obtained from PubChem and identified the top ten compounds (Limonin, Corilagin, Taraxesterol, Procyanidin, Beta Carotene, Emodin, Dehydropipernonaline, Apigenin, Guineesine, and Benzethonium) as primary candidates based on binding affinity scores with the proteins encoded by three different genotypes of *ctxB*. After doing an ADME analysis, we were able to identify three possible phytochemicals (limonin, emodin, and dehydropipernonaline) that may inhibit the cholera toxin. However, dehydropipernonaline showed hepatotoxicity in our in-silico findings.

The potential phytochemical Limonin identified in this study as an inhibitor of CT can be extracted from citrus fruits, which have been utilized as medical fruit all year round due to their various therapeutic properties, which include vitamin C, flavonoids, carotenoids, folic acid, limonoids, and high-quality soluble fibers [49,50]. Several research investigations have demonstrated the diverse biological and pharmacological properties of limonin, such as its ability to protect the liver and have antiviral, anti-inflammatory, anti-cancer, and anti-oxidative properties [51-55]. One study suggested lemon juice and vinegar as the natural substitute for antibiotics [57]. Emodin has long been utilized as an active component in herbs in traditional Chinese medicine and exhibited diuretic, antibacterial, antiulcer, anti-inflammatory, anticancer, and antinociceptive properties [58]. The fruits of *Piper retrofractum Vahl* are used for gastroprotective and cholesterol-lowering properties [59]. It contains dehydropipernonaline, which in our investigation showed anti-cholera toxin action, but it may considerably damage the liver due to its hepatotoxicity properties. Our docked complex showed that the suggested phytochemicals bind to the amino acid of *ctxB* at the several common sites as the GM1 receptor [47]. So, they can be potential compounds to inhibit *ctxB* from binding with GM1. Additionally, a molecular dynamics simulation incorporating the chosen phytochemicals and cholera toxin mutants, *ctxB1, ctxB3*, and *ctxB7*, was run to assess MM-PBSA binding free energy and RMSD of the nine complexes. All parameters consistently showed increased stability in the protein-ligand interaction, except the complex formed by emodin and *ctxB3*. Therefore, we may conclude two phytochemicals limonin and emodin to be promising for blocking cholera toxin function and preventing the deadly disease. However, validation of these results requires wet-lab synthesis and further in-vivo studies.

## Acknowledgments

This project was conducted as a non-profit collaborative effort between icddr,b, and the University of Rajshahi. icddr,b extends its gratitude to the Governments of the People’s Republic of Bangladesh and Global Affairs Canada (GAC) for their unrestricted support. We thank Kazi Ahsan from the School of Medicine at Washington University, St. Louis, USA, for his valuable comments on the draft manuscript, which helped enhance its quality.

## Author Information

### Author notes

1. Yasmin Akter and Khalilur Rahman equally contributed to this manuscript as co-first authors.

### Contributions

M.M.M, M.A., and M.N.H.M developed the study concept and facilitated the nonprofit collaborative partnership. Y.A. collected the necessary information from different sources and performed the molecular docking. The further analysis, including molecular dynamic simulation, was accomplished by K.R. Y.A., and K.R. jointly drafted the manuscript. M.S.R. contributed to organizing the method section. M.M.M. and M.N.H.M. supervised the study. All the authors reviewed and approved the final manuscript.

### Corresponding author

Correspondence to Md. Mamun Monir and Md Nurul Haque Mollah

## Data Availability

The datasets generated and analyzed during the current study will be available from the corresponding author on request.

## Competing interests

The authors declare no competing interests

## Supplementary Information

Supplementary Information

## Notes

### Competing Interest Statement

The authors have declared no competing interest.

